# Using Deep Learning to predict replication timing reveals baseline control of genomic DNA sequence

**DOI:** 10.64898/2026.07.18.739304

**Authors:** Nikola Janakievski, Pierre M. Joubert, Anna R. Poetsch

## Abstract

Human DNA replicates according to a precise schedule: certain regions are replicated early, while others replicate later in the cell cycle. The DNA sequence and the epigenome are both indicative for replication timing (RT), their relative contributions and potential complementarity yet remain unclear. Here, we utilize Repli-Seq data and machine learning to investigate these relationships in seven human cell lines. We show that GROVER (Genome Rules Obtained Via Extracted Representations), a DNA language model, trained purely on DNA sequence can be fine-tuned for RT, which indicates that information on DNA sequence is largely sufficient to predict RT. Integrating GROVER’s sequence predictions as an additional feature with diverse epigenetic sequencing data, yields a multimodal model that outperforms approaches that use exclusively the DNA sequence or the epigenome, which demonstrates predictive synergy. By evaluating input feature contributions, we find that epigenetic features vary in their RT prediction contribution. Some epigenetic features see their contribution diminish in the presence of DNA sequence information, others remain predictive. GROVER’s sequence representations emerge as the most dominant feature. Furthermore, to assess cell-type specificity of RT, we partitioned the genome into constitutive domains that replicate uniformly across cell lines and cell-type-specific domains that shift during differentiation. While DNA sequence features contribute more to constitutive predictions, they also remain informative within cell-type-specific regions. Together, our findings dissect the relationship between DNA sequence and the epigenome relative to RT and suggest that primary DNA sequence instructions exert a powerful baseline control over constitutive and cell-type-specific RT, fine-tuned by regulatory cues beyond the DNA sequence.

## Introduction

The human genome replicates according to a precise schedule distinguishing sites for replicating early or late in the cell cycle. The replication timing (RT) program impacts replication stress and partitioning mutational loads across the genome (1–3). It is largely conserved across individual cells of the same type (4,5) but remains highly dynamic during differentiation (6,7) and cancer development (8–10). Replication timing correlates strongly with various epigenetic marks and has been proposed as a master regulator of the global epigenetic state (11). Early RT is correlated with high chromatin accessibility, gene density and to a lesser extent by transcriptional output (6,7,12). Specifically, active histone modifications such as H3K4me3 at promoters, H3K27ac at promoters and enhancers, and H3K36me3 within gene bodies are enriched in early-replicating domains (6,13,14). The constitutive heterochromatin mark H3K9me3 is found within late-replicating domains (13). The architectural protein CCCTC-Binding factor (CTCF) and the facultative heterochromatin mark H3K27me3 appear in timing transition regions (14). Based on these associations, predictors of RT have been successfully developed on epigenetic data (15,16).

The epigenetic landscape is closely linked to, and often driven by the underlying genomic sequence (17,18), which applies to RT regulation as well. Early replicating control elements (ERCEs) are enriched in various transcription factor binding sites, including for CTCF, and active histone marks (19,20). Asynchronous replication and autosomal RNAs (ASARs) are DNA sequences that encode long non-coding RNAs which regulate RT. (21). Beyond cis-acting elements, sequence features like GC-content, non-B-DNA structures like G-quadruplexes, and repetitive elements have long been known to correlate with RT (6,7,22), and have been leveraged to build models to predict RT. With the context-aware model Concert, the DNA sequence patterns alone contain sufficient instructional signals to accurately predict RT profiles while simultaneously identifying key underlying motifs that drive RT (23). Predictive performance varies across different cell lines and genomic regions. Recently, the Soffritto model (24) employed a Long Short-Term Memory (LSTM) based architecture to enhance Repli-seq data resolution by integrating sequence and epigenetic data.

Foundational DNA language models (LLMs) like GROVER (Genome Rules Obtained Via Extracted Representations) utilize unsupervised pretraining across the entire genome to learn a universal, contextual DNA syntax before any task-specific fine-tuning occurs (25). During this self-supervised phase, GROVER implicitly captures broad genomic architectures and functional features without any explicit labels, including GC content, gene loci, repetitive elements, and critically, RT itself. However, genomic function is rarely defined by DNA alone; it emerges from the complex interplay between the sequence and the chromatin environment (26–28).

We recently employed a multimodal learning strategy leveraging GROVER to explore the relative contributions of DNA sequence and chromatin context in modeling genomic double-strand break (DSB) sensitivity (29). We now build on this framework to examine the sequence dependence and cell-type specificity of RT to study the contribution of DNA sequence information and the epigenome in predicting RT across different cell types using GROVER as a proxy for the relative contribution of DNA sequence, integrated via a random forest model with epigenetic data. We reveal a synergistic relationship that substantially boosts predictive performance. Using model interpretability methods, we find that the importance of some epigenetic features diminishes when DNA sequence information is included, while others retain their predictive power, with GROVER sequence representations consistently emerging as the dominant contributor. Decomposition of these contributions shows that sequence is the primary driver of constitutive domains and remains a critical baseline for RT regulation even within cell-type-specific regions, while some epigenetic features contribute more to constitutive and others more in cell-type specific RT.

## Results

### GROVER can largely predict replication timing based on DNA sequence alone

The DNA sequence is a strong determinant of replication timing (7,19,22,30). To investigate the degree to which RT is encoded in the DNA sequence, we fine-tuned the foundational DNA language model GROVER (25) to predict RT across seven ENCODE top-tier cell lines (GM12878, K562, HeLaS3, HepG2, HUVEC, MCF7 and NHEK) (18) (**Fig. 1A**). RT scores were derived from Repli-seq data for genomic bins matching the model’s maximum input size (2,235 bp on average). Given that RT is a domain-level feature spanning hundreds of thousands of base pairs (3), we increased the model’s range by utilizing the learned sequence representations from fine-tuned GROVER as input to a LSTM network with an input size of 100-bin windows (∼223 kb), the saturation point of context size (**Supp. Fig. 1**). GROVER predicts RT across seven cell lines with Spearman correlation coefficients between 0.747 in HeLaS3 and 0.835 in K562 (**Fig. 1B**) and predictions that recapitulate the characteristic bimodal distribution of RT measurements (**Fig. 1C**). GROVER outperforms the model Concert in K562 (Sp. R = 0.835 versus 0.783) and performs on-par in GM12878 (Sp.R = 0.831 versus 0.838) (23). GROVER leverages its pre-learned knowledge to map DNA sequence features to the RT profiles of different cells. This demonstrates that sequence patterns alone encode substantial information on RT. Longer-range sequence patterns are particularly relevant, because the same task for GROVER without the LSTM leads to inferior performance (Sp. R = 0.544 to 0.660; **Supp. Fig. 2A**). GC-content is a known determinant of RT (6,7,22). We find that while GC-content correlates with RT (Sp. R = 0.511) (**Fig. 1D**), the correlation of the GROVER’s prediction performance with RT exceeds the contribution of GC content alone (Sp. R = 0.835). This also varies among cell lines. Cells whose RT programs exhibit the strongest correlation with GC content are also the most accurately predicted by GROVER (**Supp. Fig. 3)**. Interestingly, GROVER’s prediction outputs show a stronger correlation with GC-content than Repli-seq measurements (Sp. R = 0.583 vs. 0.511, respectively). This could be explained by GROVER overly relying on DNA sequence, even beyond the RT being affected by it, given the sequence-independent determinants that are indicated by the variability in cell-type specific regions (**Fig. 1E**). Since all cell types in an organism share a nearly identical genome, a sequence-only model is inherently unable to detect regions that shift replication timing across cell types, which suggests that integrating DNA sequence and epigenetic data into a multimodal model of RT should improve RT prediction even further.

**Figure 1.**
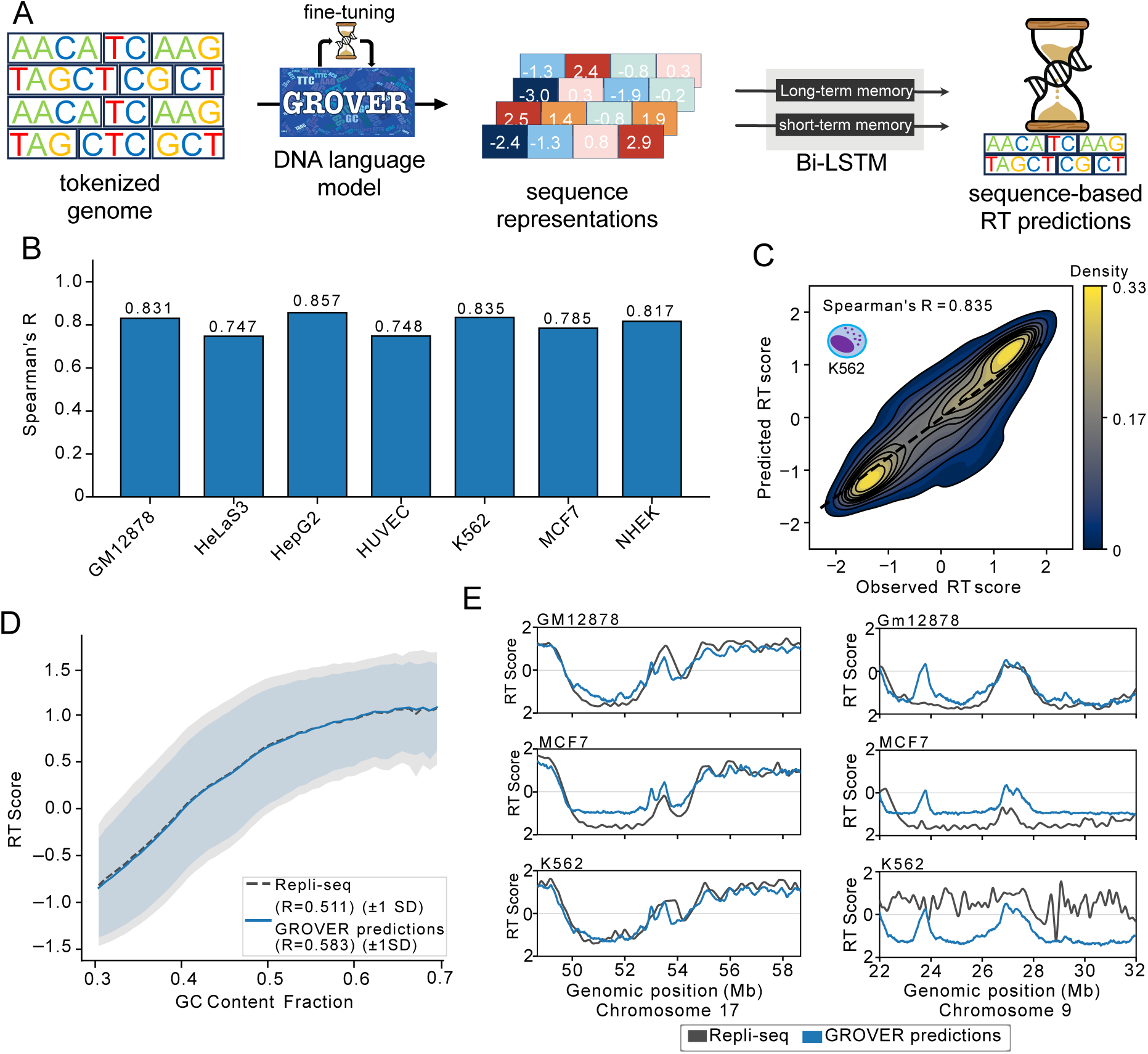
GROVER can largely predict replication timing (RT) based on DNA sequence alone. **A.** In the sequence model, GROVER embeddings fine-tuned specifically for RT prediction are used to train a Long Short-Term Memory (LSTM) network, which expands context across 100 contiguous genomic bins to predict RT from sequence alone. **B.** Performance of the sequence model across seven cell types. **C.** Distributions of observed and sequence-model predicted RT for a representative cell line. **D.** The relationship between the GC-content of a genomic bin and the observed and DNA sequence-model predicted RT scores. **E.** Comparison of the observed and DNA sequence-model predicted RT scores across three different cell types in two different genomic regions.

**Figure 2.**
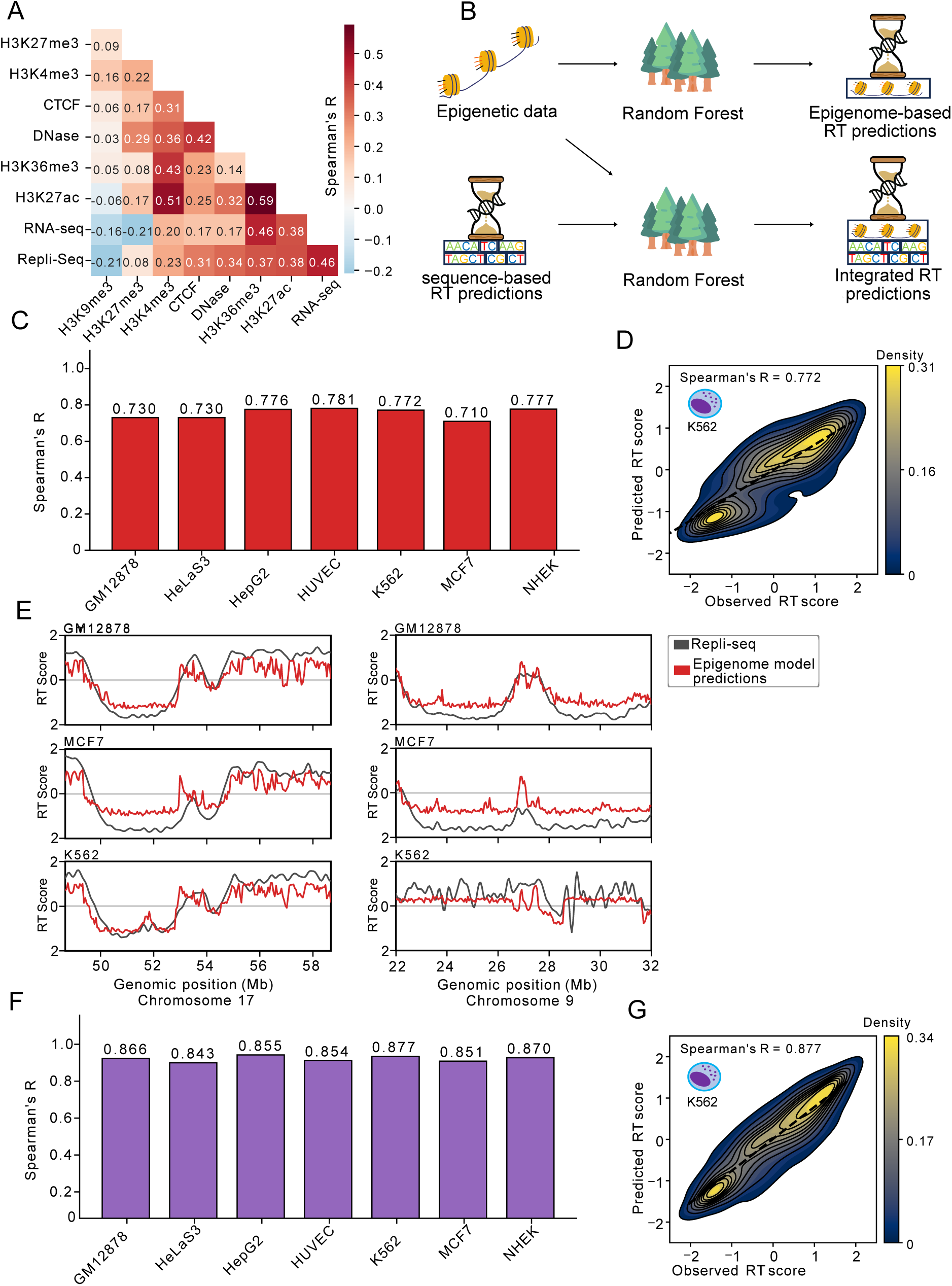
DNA sequence and epigenetic context provide complementary information for predicting replication timing (RT). **A.** Spearman’s correlation coefficients between various epigenetic features and Repli-seq signals, averaged across seven cell lines. **B.** In the epigenome and integrated models, random forests are trained on either epigenetic features alone or both epigenetic features and DNA sequence-model predictions to predict RT. **C.** Performance of seven cell-type epigenome models. **D.** Distributions of observed and epigenome-model predicted RT for a representative cell-type model. **E.** Comparison of the observed and epigenome-model predicted RT scores across three different cell types in two different genomic regions. **F.** Performance of seven cell-type integrated models. **G.** Distributions of observed and integrated-model predicted RT for a representative cell-type model.

**Figure 3.**
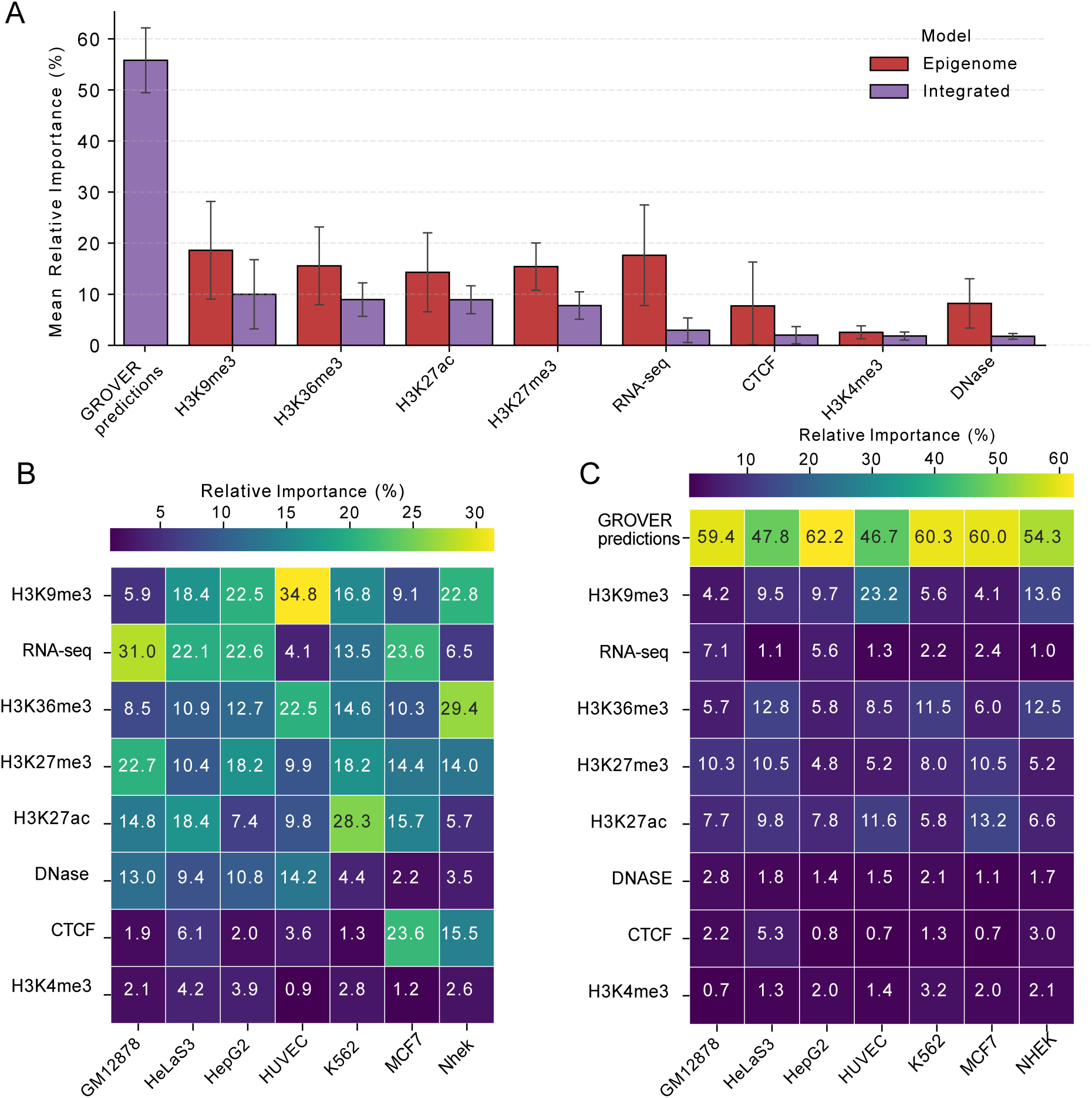
GROVER predictions are the most important feature for predicting replication timing (RT). **A.** Relative importance of features used by the epigenome and integrated models to predict RT, averaged across seven distinct cell-type models. Relative importance is calculated by dividing a feature’s mean absolute SHAP value by the sum of the mean absolute SHAP values of all features in that specific model. **B.** Relative importance of features for the seven epigenome cell-type models. **C.** Relative importance of features for the seven integrated cell-type models.

### DNA sequence and epigenetic context provide complementary information for predicting replication timing

To investigate the relationship of RT with epigenetic data, we built a model trained exclusively on epigenetic information from various ENCODE sequencing datasets (18) for the same seven cell lines. Consistent with previous studies (6,7,13,14,31), we observe that RT correlates with various epigenetic features (**Fig. 2A**). H3K9me3 was the sole epigenetic feature to negatively correlate with Repli-seq (Sp. R = -0.21), reflecting its enrichment in late-replicating domains (6). H3K27me3 showed a comparatively lower correlation (Sp. R = 0.08), likely due to its prevalence in timing transition regions (TTRs) (14). Despite its link to TTRs (14), CTCF protein binding correlates more strongly with Repli-seq (Sp. R = 0.31), potentially driven by its role in early replicating control elements (ERCEs) (19). There are also strong correlations between the epigenetic features, indicating that they provide redundant information for predicting RT.

To model RT with epigenetic features, we followed a modelling approach previously used for genome instability (29,32). In brief, we trained a random forest model per cell type to predict RT on the same genomic bins used for GROVER (**Fig. 2B**). The epigenome models achieve Spearman’s correlations ranging from 0.710 to 0.781 (**Fig. 2C, 2D**). While the absolute performance of the epigenome models is lower than the sequence model, they are able to capture cell-type specific RT (**Fig. 2E**).

The epigenome has strong correlation with sequence features and can to a large extent also be predicted based on the DNA sequence, as shown by the recent developments with AlphaGenome (28). However, integrating both sequence and epigenome information leads to improved predictions, when using the sequence model’s predictions as an additional training feature within the epigenome models (Sp. R = 0.843 – 0.877) (**Fig. 2F, 2G**) and yields high stability across cell lines. This suggests that the GROVER-learned sequence representations and the epigenome context provide synergistic information for predicting RT.

### DNA sequence is the most important single feature for predicting replication timing

To further investigate the synergy between the DNA sequence and epigenetic features, we quantified the contribution of each feature to the epigenome and integrated predictions using SHapley Additive exPlanations (SHAP) (33) (**Fig. 3A**). We found that feature importances do not strictly follow the previously observed feature correlations to RT (**Fig. 2A**). RNA-seq and DNase-seq are both strongly correlated and very predictive, confirming the link between transcriptional output, chromatin accessibility and early replication (6,7). H3K9me3 and H3K27me3 are some of the most important features for the epigenome models despite modest correlations (Sp. R = -0.21 and 0.08, respectively). This reflects H3K9me3’s unique role as the primary marker for late-replicating domains (13) and the H3K27me3 in TTRs (14). Features with strong correlations to RT, such as H3K36me3, H3K27ac, H3K4me3 associated with enhancers, transcriptional elongation and promoters respectively (34–36), are all found in early-replicating regions (6) and as a consequence provide overlapping information, possibly resulting in the relative low importance of H3K4me3. GROVER’s predictions are the single most important feature for the integrated models with a mean relative importance of 55.61% (**Fig. 3A**). While larger context sizes enhance sequence model performance (**Supp. Fig. 1**); (23), GROVER’s dominance persists even when controlling for this variable. Specifically, predictions from a model limited to 2.2 kb remained the most influential feature at 34.08% (**Supp. Fig. 2E**), confirming that sequence dominance is not an artifact of context size. Comparing feature importances between the integrated and epigenome models revealed that some features (**Fig. 3A**), like RNA-seq, DNase-seq, and CTCF binding sharply decline in importance upon addition of sequence information, while the histone marks remain stable. As an example, CTCF binding can be predicted by GROVER (25), meaning GROVER’s sequence representations already contain much of this information. This result indicates that while DNA sequence and the epigenome share some overlapping information, the degree of redundancy is feature-specific.

Feature importances within different cell-type epigenome models vary (**Fig. 3B**). This is relatively minor for DNase-seq and H3K4me3, but very pronounced for all other features. For example, H3K9me3 is the most important feature for the model for HUVEC cells, but only the 6th most important for the GM12878 model. Both differential feature contributions and model performance could be due to biological differences between the cell lines, yet could also be affected by data quality. While the sequence remains the most important feature in integrated models across cell lines, its relative importance varies (**Fig. 3C**). The cell lines where the DNA sequence models performs worst are also the ones that rely least on sequence-based predictions (**Supp. Fig. 3**).

### Model predictive performance diverges between constitutive and cell-type specific replication timing

To evaluate how cell-type specificity influences model performance, we annotated the genome for constitutive and cell-type specific bins based on RT variability across the seven cell lines (**Fig. 4A**). The variance threshold divides the genome into halves to align with previous estimates that approximately 50% of the genome undergoes RT shifts during differentiation (7,37,38). Consistent with previous studies, we find that constitutive regions generally replicate earlier (**Fig. 4B**) and are significantly enriched in gene density and regulatory elements (6,7,38) (Fisher’s exact test, p < 0.001; **Fig. 4C**). Fine-tuned GROVER accurately captures constitutive RT (Sp. R = 0.865 to 0.900) (**Fig. 4D, 4G**). In cell-type specific regions, although accuracy decreases (**Fig. 4D, 4J**), the model nonetheless captures RT signal (Sp. R = 0.432 to 0.691), which suggests that DNA sequence remains a relevant predictor of RT in these domains. The epigenome models showed more balanced performance (**Fig. 4E, 4H, 4K**), though cell-type specific bins remain harder to predict (Sp. R = 0.577 to 0.713) than constitutive ones (Sp. R = 0.705 to 0.770). This disparity likely arises from the enrichment of genes and regulatory DNA in constitutive versus cell-type specific bins, which provides a denser information landscape. The relative sparsity of cell-type specific bins reduces the effective signal available to the epigenome models, making it more challenging to generalize patterns in these regions. By combining sequence and epigenome context, the integrated models are able to match the sequence model’s accuracy in constitutive regions (Sp. R = 0.875 to 0.902) while greatly improving on cell-type specific bins (Sp. R = 0.686 to 0.808) (**Fig. 4F, 4I, 4L**). That sequence remains informative in cell-type specific regions challenges the assumption that constitutive RT regions would be more dependent on DNA sequence.

**Figure 4.**
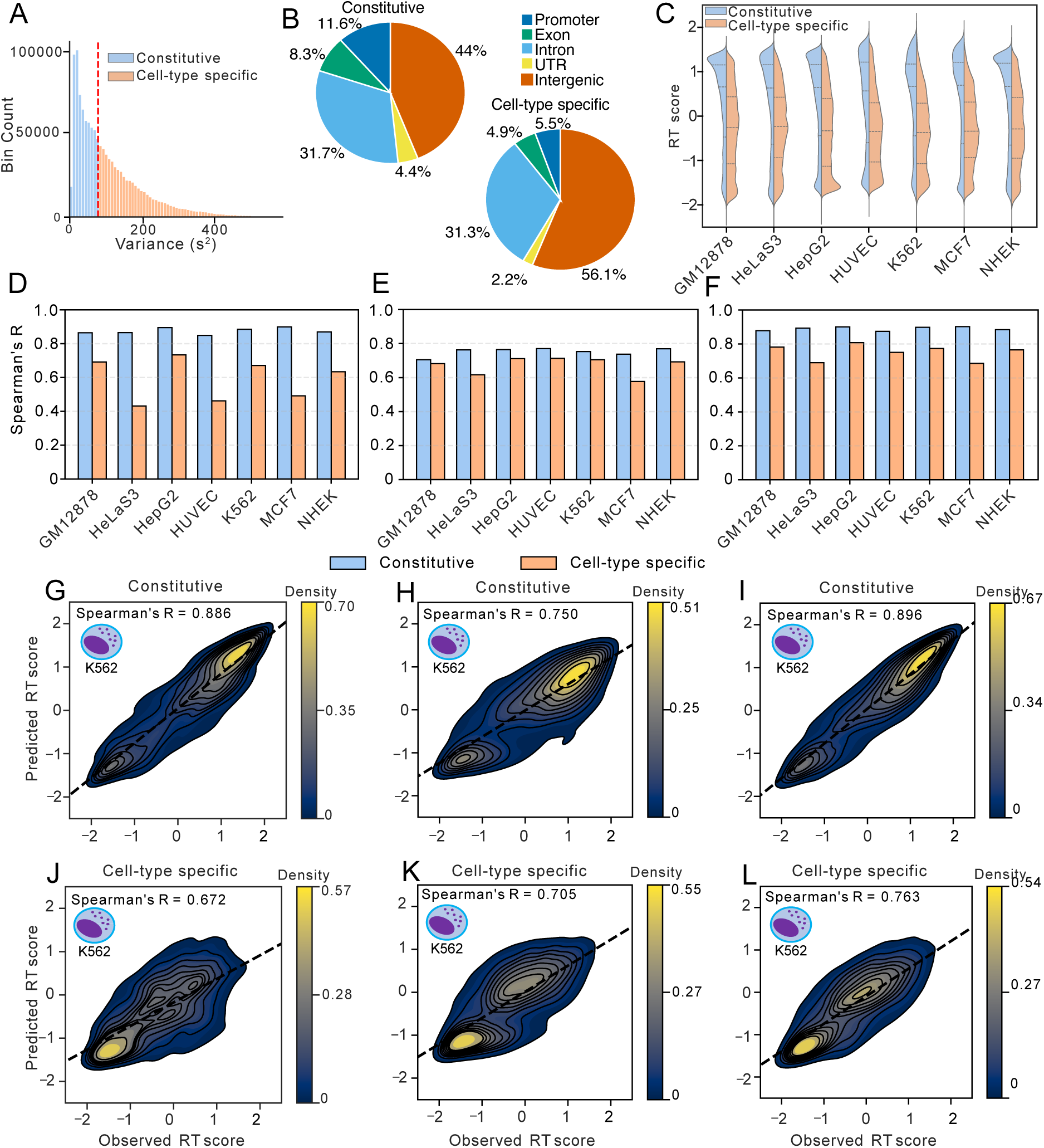
Model predictive performance diverges between constitutive and cell-type specific replication timing (RT). **A.** Distribution of RT score variance across the seven cell-type models. Genomic bins are classified as constitutive or cell-type-specific based on a median variance threshold. **B.** Genomic distribution of annotations (promoters, exons, introns, UTRs, and intergenic regions) within constitutive and cell-type-specific bins. **C.** Distribution of RT scores across constitutive and cell-type-specific bins for each of the seven cell lines. **D, E and F.** Performance of the DNA sequence model, and the seven individual epigenome and integrated cell-type models, across all seven cell lines predicting RT scores in constitutive and cell-type-specific bins. **G, H and I.** Distributions of observed and predicted RT scores within constitutive bins for a representative cell type, shown for the DNA sequence, epigenome, and integrated models, respectively. **J, K and L.** Distributions of observed and predicted RT scores within cell-type-specific bins for a representative cell type, shown for the DNA sequence, epigenome, and integrated models, respectively.

### DNA sequence information contributes more to constitutive than to cell-type specific predictions

To further investigate the disparity between DNA sequence and epigenome contributions, we evaluated the reliance of RT on GC-content in both constitutive and cell-type specific domains (**Fig. 5A, 5B**). We found that constitutive RT correlates more strongly with GC-content than cell-type specific RT (Sp. R = 0.554 versus 0.380). The DNA sequence model partially recapitulates this trend, as its RT predictions correlate less with GC-content in cell-type specific domains compared to constitutive ones (Sp. R = 0.508 versus 0.595). To complement this analysis, we stratified feature importances across either strictly cell-type specific or constitutive genomic bins (**Fig. 5C, 5D**). We observed that some features such as RNA-seq (+1.40% relative importance), Dnase-seq (+1.10%) and H3K36me3 (+1.98%), contribute more to the prediction of constitutive RT. Conversely, repressive histone marks, H3K27me3 (-2.72%) and H3K9me3 (-1.68%), exhibit higher relative importance in predicting cell-type-specific RT compared to constitutive regions. In our integrated model, these patterns persist, with repressive marks remaining more influential in cell-type-specific regions (e.g., -3.39% for H3K9me3 and -2.50% for H3K27me3 relative to constitutive bins). Most importantly, we find that GROVER predictions contribute more to constitutive than to cell-type specific RT (59.7% and 51.4%, respectively), indicating that DNA sequence information remains deeply informative of RT even within cell-type specific domains.

**Figure 5.**
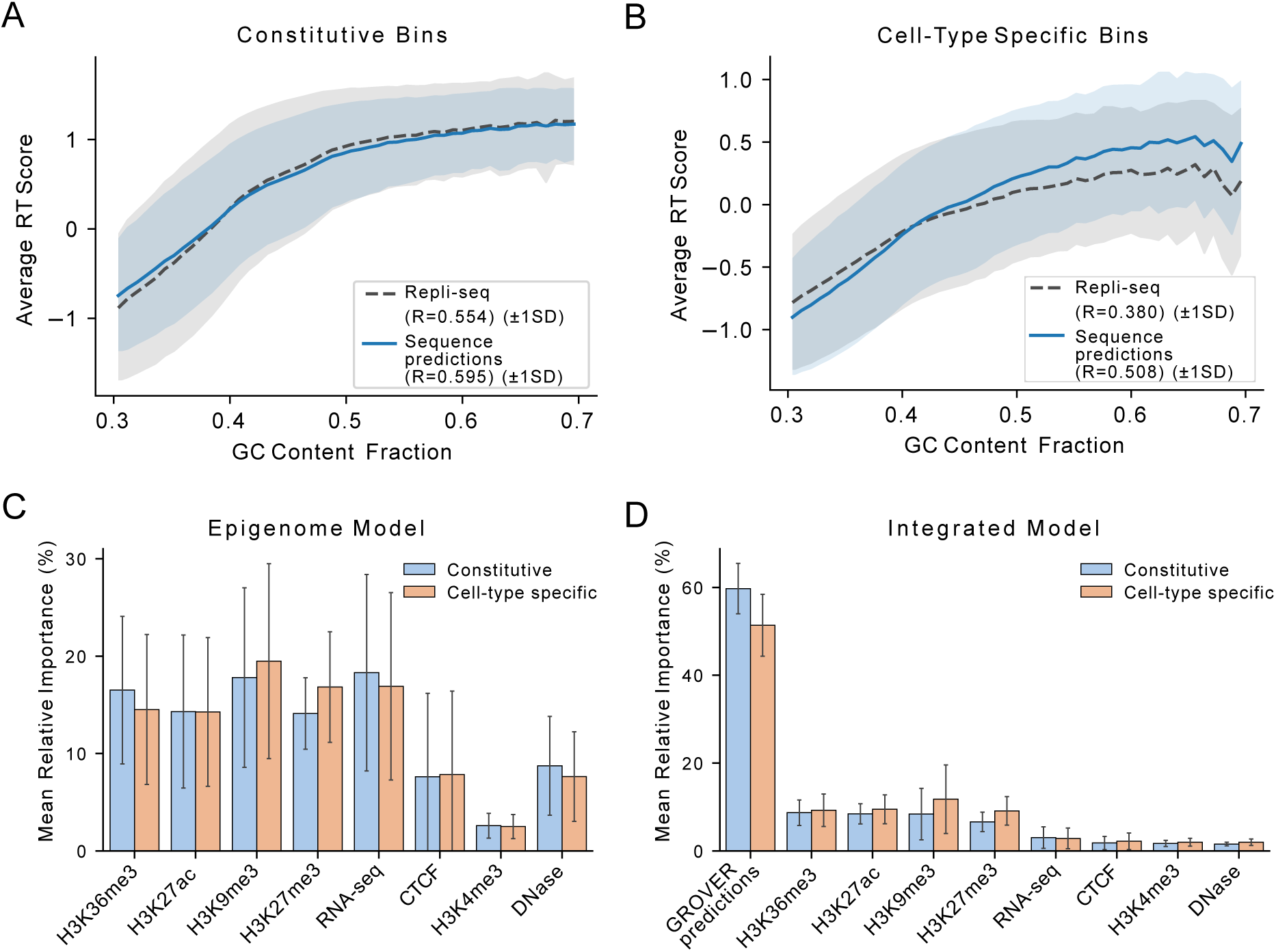
GROVER predictions contribute more to constitutive than to cell-type-specific replication timing (RT) predictions. A and. **B.** The relationship between the GC-content of a genomic bin and the observed and DNA sequence-model predicted RT scores for constitutive and cell-type-specific bins, respectively. **C and D.** Relative importance of features used by the epigenome and integrated models, respectively, to predict constitutive and cell-type-specific RT, averaged across seven distinct cell-type models.

## Discussion

By fine-tuning GROVER to predict RT, we confirmed previous findings that RT can be modeled from DNA sequence alone (23,39). The model’s ability to predict RT is primarily guided by the RT dependence on GC-content, aided by the recognition of sequence motifs such as CTCF and transcription factor binding sites found in ERCEs (19). GROVER’s RT predictions could further be informed by the density of sequence features near replication origins (40), such as G-quadruplexes and CpG islands. Finally, GROVER’s detection of retrotransposon sequences (25) might improve its RT predictions by identifying ASARs, whose disruption leads to delayed replication on their native chromosomes (21). Fine-tuning across seven different cell lines also allowed GROVER to learn the relationship between DNA sequence and different cellular context and allowed for the model to associate specific sequence patterns with context-dependent RT outcomes. Yet, interestingly, performance varies greatly across cell lines. Because the DNA sequence inputs are identical, these discrepancies can stem from three factors: variations in Repli-seq data quality, genomic divergence from the human reference genome, particularly in unstable, rearranged lines like K562 and HepG2 (41,42), where sequence alterations can shift RT profiles (30,43), or varying degrees of cell-type-specific epigenetic influence that alter the contribution of sequence as a predictor of RT.

Indeed, including epigenetic context both improves and homogenizes model performance across cell types. This synergistic interaction between DNA sequence and epigenome is increasingly recognized in genomic modeling. Studies on gene expression (44,45), DNA methylation (46), and DSBs (29) demonstrate that sequence-only models reach a plateau that is overcome by incorporating cell-type specific epigenetic information. Experimental functional assays also support this view, demonstrating that a DNA sequence’s regulatory activity is not independent, but is influenced by its epigenetic context (47,48). This finding parallels existing models of RT control, where ERCEs have been characterized as master regulatory hubs (19,20). Within these elements, sequence motifs provide the foundation, but their activity is contingent on cell-type specific transcription factor binding and histone marks. While previous studies have established that DNA sequence (23,24,39) and the epigenome (15,16) are both predictive of RT, their relative importance remained poorly defined. In our integrated model, DNA sequence emerges as the dominant contributor for predicting RT. This contrasts with studies on gene expression (49) and DSBs (29), where sequence is one of several influential features, which suggests that RT is more dependent on the DNA sequence. This high sequence dependence of RT could be a result of its proposed role as a maintainer of the epigenetic state (11,20,50). The addition of DNA sequence information diminished the importance of RNA-seq, Dnase-seq, and CTCF for RT prediction. This redundancy arises because these features are largely DNA sequence-intrinsic: RNA-seq is a readout of transcriptional (coding) potential (51), while DNase-seq and CTCF binding are governed by specific sequence motifs (17), the latter can be accurately predicted by GROVER (25). However histone marks all remain robust, suggesting that histone marks provide more complementary information to sequence. This is likely because they serve as predictors of chromatin state (52) and capture classes of epigenetic function across cell lines that are not fully encoded in the DNA sequence. RT shifts have been shown to influence histone acetylation patterns (19,53). H3K9me3 and H3K36me3 levels have been linked to RT regulation by H3K9/36 tri-demethylase KDM4A overexpression (31,54). H3K27me3 is enriched in TTRs and acts as a determinant for intermediary RT states (14,31). Besides enrichment in early RT (13), H3K4me3 has not been found to directly interact with RT and that could explain its low importance. These findings underscore that while DNA sequence is the dominant predictor of RT, the collective regulation of histone modifications provides the cellular context to capture the full dynamics of the RT.

Our variance-based annotation of the genome into constitutive and cell-type-specific RT regions aligns with established literature showing that constitutive regions are typically gene-rich and replicate early, while cell-type specific regions replicate later and consist of more intergenic DNA (6,7,38). This dichotomy explains the overall lower model performance in cell-type specific bins. The reduced density of gene and regulatory features limits the predictive signal available to both sequence- and epigenome-based models. This also explains the increased feature importance in cell-type specific RT prediction of inaccessibility-associated histone marks (H3K27me3 and H3K9me3). GROVER’s predictions are expectedly more important for constitutive RT, but surprisingly they remain robust in cell-type specific RT. This indicates that cell-type-specific RT programs are not purely epigenetic in origin but are inherent in the DNA sequence. The distinct sequence composition of cell-type specific RT regions points to this (38), which could also inform GROVER’s predictions. The overall high importance of sequence information for RT prediction revealed in this study, aligns with the evolving perspective on RT regulation. While early understanding of RT emphasized the role of epigenetic context (6,13,37), more recent studies have begun to integrate the contribution of DNA sequence (19,21,30). Indeed, DNA sequence-encoded instructions exert a baseline control over RT that persist even within cell-type-specific genomic regions.

Our study advances the understanding of how DNA sequence and epigenetic context act in concert to shape both constitutive and cell-type-specific RT. By leveraging deep learning as an analytical lens, we demonstrate that these factors can be disentangled to reveal their relative contributions to the genome’s functional program. We reveal that DNA sequence as a baseline persists even within cell-type-specific RT. While our focus was the RT program, this multimodal interpretability strategy is applicable to other genome functions that are governed by complex regulatory principles, such as 3D genome organization or genome instability.

## Acknowledgements

The authors express their gratitude to the NHR Center at TU Dresden for providing the high-performance computing resources necessary for this work. This center is jointly supported by the Federal Ministry of Education and Research and the state governments participating in the NHR (www.nhr-verein.de/unsere-partner). We thank Melissa Sanabria for her help with implementing the GROVER model. Additionally, we appreciate Mario Aguilar-Herrador’s valuable insights regarding data visualization and thank the Biomedical Genomics group at TU Dresden for their ongoing support and insightful discussions.

## Methods

Unless stated differently, Machine learning and data preparation scripts were written in Python (v3.11.11) with Scikit-Learn (v1.7.1), NumPy (v2.2.6), PyTorch (v2.5.1+cu121), Pandas (v2.3.3), SHAP (v0.51.0), SciPy (v1.16.1) and Transformers (v4.57.1). Visualizations were created using Matplotlib (v3.10.6) and Seaborn (0.13.2) with some plots created via PyBigWig (v0.3.25) and ChIPseeker package in R (v4.3.1) as described in their respective sections.

### Replication timing data

Repli-seq data for the cell lines was acquired from the ENCODE project (18) (UCSC_Track_Identifiers.xlsx). A yet unpublished 1000 cycles of byte-pair encoding GROVER-tokenized *Homo sapiens* (human) hg19 (GRCh37) genome was divided randomly into non-overlapping bins of 510 tokens (on average 2235 bp), with each bin serving as a discrete input sequence for model training or prediction. Any bins containing Ns, overlapping with centromeric, telomeric regions or the ENCODE blacklist (55) were removed from the dataset. The average WaveSignal value over each genomic bin was calculated using bigWigAverageOverBed (56).

### GROVER fine-tuning

We fine-tuned GROVER to simutaneously predict RT score of seven different cell lines. To generate genome-wide sequence representations, we employed a 5-fold cross-validation strategy. Within each fold, the genome was partitioned into training (70%), validation (10%), and test (20%) sets. Outlying target values (>99.9%) and zeroes were removed. The data were then z-score normalized using StandardScaler from scikit-learn (v1.7.1), with parameters calculated exclusively from the training set. The pretrained GROVER model (unpublished) was loaded using transformers (v4.57.1) and fine-tuned using torch (v2.5.1+cu121). Fine-tuning was done with a Mean Squared Error (MSE) loss function, using the AdamW optimizer with an epsilon of 10^−8^, a learning rate of 10^−6^, warm up of 1000 steps, a batch size of 128, and a dropout probability of 0.3. Each k-fold iteration was fine-tuned for 5 epochs and only the best performing model was kept, based on evaluation on the validation set after every epoch. Afterwards, the mean pooled embeddings for each genomic bin from each test set were extracted and concatenated.

### LSTM training

To predict replication timing, we trained a two-layer bidirectional LSTM (1024 hidden states) using the extracted GROVER embeddings (768-dimensional). The model processes sequences of 100 consecutive bins (context size chosen based on optimal performance; **Supp. Fig. 1**) to generate one prediction per bin across seven cell lines. Training and evaluation were conducted via 5-fold cross-validation (70/10/20 split), with a mask applied during loss calculation to ensure the model was optimized and evaluated exclusively on genomic bins that have Repli-seq measurements. Target values were standardized using StandardScaler (scikit-learn v1.7.1) and fitted exclusively on the training subset of each respective fold. Early stopping was implemented based on validation loss to select the optimal model state for each fold. The model was implemented using PyTorch (v2.5.1+cu121). The predictions of each test set were concatenated together to yield genome-wide predictions, which are then used as a feature when training the integrated model. Model performance was evaluated on the unseen test set across all, cell-type specific, and constitutive genomic bins with the best performing fold.

### Epigenetic data

All of the epigenetic data (RNA-seq, DNase-seq, CTCF ChIP-seq, H3K27me3, H3K27ac, H3K4me3, H3K9me3 and H3K36me3 ChIP-seq) for all seven cell lines used to train the epigenome and integrated model were obtained from the ENCODE project (The ENCODE Project Consortium 2012) (UCSC_Track_Identifiers.xlsx). To generate a features table, we mapped epigenomic feature tracks (BigWig) to the genomic bins using bigWigAverageOverBed (v4) (56), which calculates the average signal for each bin.

### Epigenome Model

Random forest models (500 trees, max depth 15) were trained per cell type using scikit-learn (v1.7.1). To maintain consistency with the GROVER model, we utilized the same genomic binning: each model was trained on data derived from the same 510-token bins (averaging 2,235 base pairs). Genomic bins were partitioned into training, validation, and test sets (80/10/10 split), with 10 independent iterations performed to ensure robustness. After removing zeroes and outliers (>99.9%), target values were z-score normalized using StandardScaler from scikit-learn (v1.7.1) fitted exclusively on the training set. Model performance was evaluated on the unseen test set across all, cell-type specific and constitutive genomic bins.

### Integrated model

To integrate sequence-based predictions into the epigenetic model, we reconstructed a genome-wide dataset from the five mutually exclusive LSTM test folds. These predictions were concatenated with the epigenetic feature table to train random forest models. Performance was evaluated on the unseen test set across all, cell-type specific and constitutive genomic bins.

### SHapley Additive eXplanations

Feature importance was determined using TreeExplainer from the shap library (v0.51.0) (33). SHAP values were computed for a random 10% subsample of the test set, with results partitioned into constitutive or cell-type specific categories. The mean absolute SHAP value per feature was calculated and then normalized by the sum of mean absolute SHAP value of all features within an individual model to derive relative importance across all, cell-type specific and constitutive genomic bins.

### Definition of cell-type specificity

To determine the cell-type specificity of a genomic bin, we calculated the variance of RT score across all seven cell lines for each bin. Bins were categorized as cell-type specific or constitutive based on a median-split threshold of the calculated variance.

### Whole genome replication timing predictions genomic tracks

To generate genome-wide RT tracks, the LSTM and RF models were trained via 5-fold cross-validation (70/10/20 split), then the predictions of each test set were concatenated to generate genome-wide predictions. These model predictions were then mapped to genomic coordinates and formatted as bedGraph files. These files were sorted by chromosomal position and converted to BigWig format using the bedGraphToBigWig utility (56) and hg19 chromosome sizes. To maintain track continuity, bins with missing predictions were assigned a fill value of 0.0.

### Genome annotation

Genomic bins were annotated using the ChIPseeker package in R (v4.3.1) (57,58), based on UCSC hg19 knownGene (59). Genomic bins were annotated following a hierarchy. Promoters were assigned to bins within 3 kb of a transcription start site (TSS). Remaining bins were assigned to exons, introns, or untranslated Regions (UTRs) in that order. To simplify the final distribution, downstream and distal intergenic regions were consolidated into a single intergenic category. Fisher’s exact test from scipy (v1.16.1) was used to evaluate if the differences in annotation distribution were significant between constitutive and cell-type specific bins.

### Density plots

Model performance was visualized using kernel density scatter plots. Density was estimated using a Gaussian kernel with 20 contour levels (ranging from 0.05 to 1.0). Plots represent a random 10% subsample of the test set.

## Declarations

### Ethics approval and consent to participate

Not applicable.

### Availability of data and materials

All code, models, input and analysis files are available on Zenodo (doi:10.5281/zenodo.21414391).

### Competing interests

The authors declare that they have no competing interests.

### Funding

This work was supported by the Center for Scalable data analytics and artificial intelligence (Scads.AI) Dresden-Leipzig. This work was partially funded by the Center for Advanced Systems Understanding (CASUS) which is financed by Germany’s Federal Ministry of Education and Research (BMBF) and by the Saxon Ministry for Science, Culture, and Tourism (SMWK) with tax funds on the basis of the budget approved by the Saxon State Parliament. **ARP** was supported by the Mildred Scheel Early Career Center Dresden P2, funded by the German Cancer Aid. **NJ** was supported by the TU Dresden programme ‘FOSTER – Funds for Student Research’.

### Author contributions

**ARP** conceptualized the study with help from **PMJ**. **NJ** and **PMJ** trained the models. **NJ** and

**ARP** analyzed the data. **NJ** and **ARP** wrote the manuscript.

---

**Figure S1.**
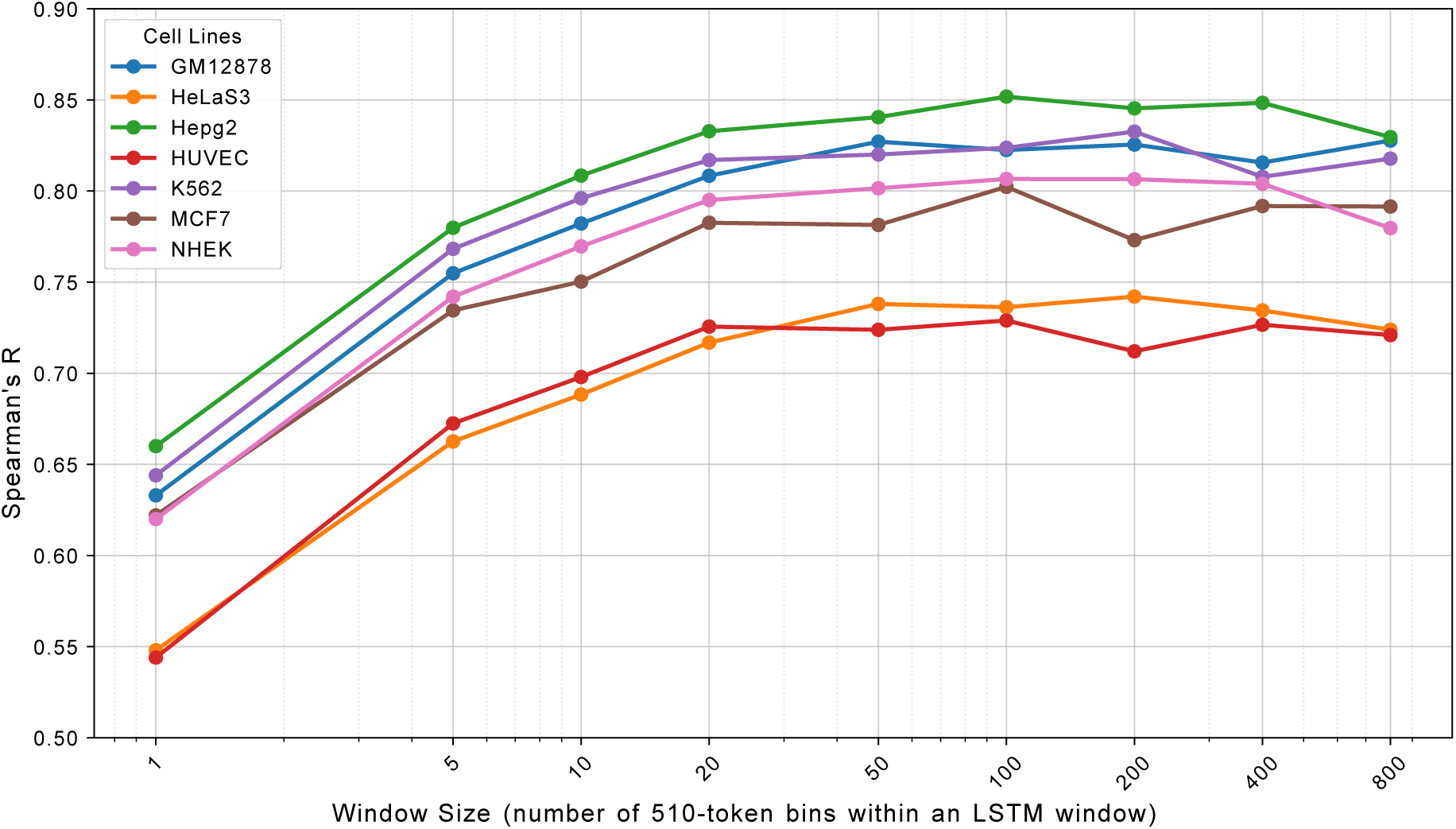
The relationship between sequence model performance and input sequence length. LSTM models were trained on different context sizes and their performances was compared. 510 tokens are on average 2235 base pairs long.

**Figure S2.**
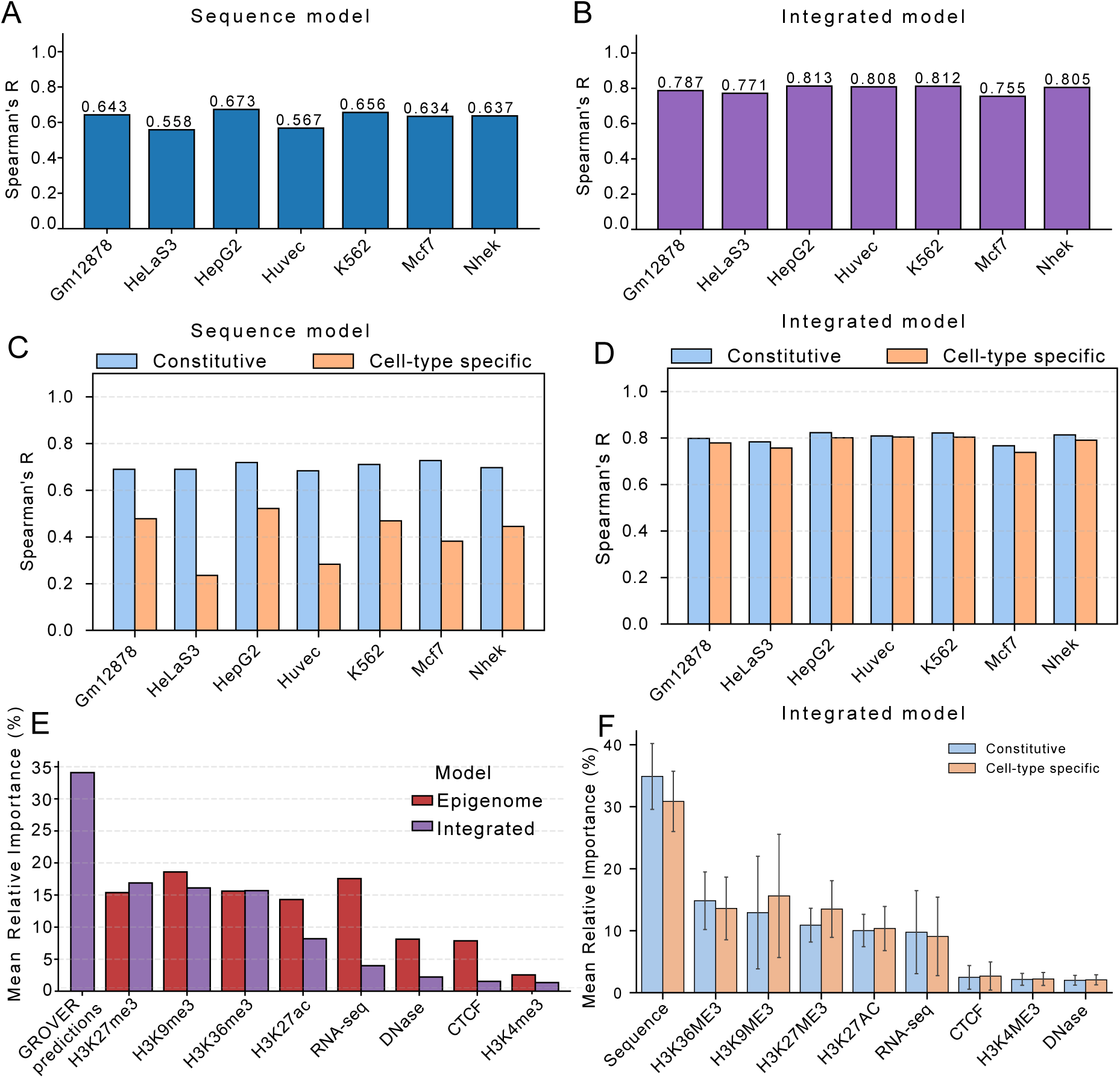
Major results redone without the LSTM. **A.** Performance of base fine-tuned GROVER (average input sequence length of 2,235 base pairs) across seven cell types. **B.** Performance of seven cell-type integrated models (base GROVER predictions). **C and D.** Performance of base fine-tuned GROVER and the seven integrated cell-type models across all seven cell lines predicting RT scores in constitutive and cell-type-specific bins. **E.** Mean relative importance of features used by the epigenome and integrated models **F.** Mean relative importance of features in the integrated model used to predict constitutive and cell-type-specific RT

**Figure S3.**
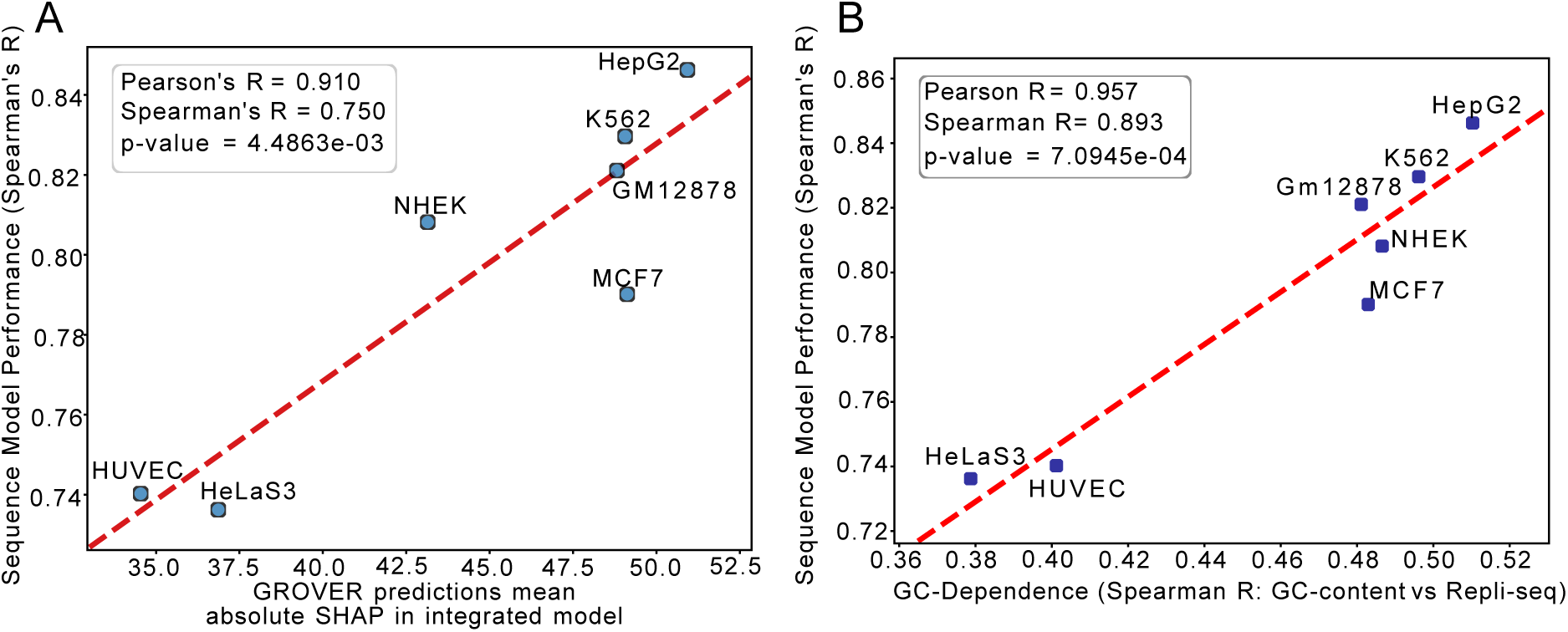
Sequence model performance correlates with GC-dependence and sequence importance. **A.** Significant correlation between the sequence model’s performance and the GROVER predictions’ contribution in the different cell-type integrated models. **B.** Significant correlation between the sequence models’ performance and the GC-dependence of Repli-seq of different cell types.

## Notes

https://doi.org/10.5281/zenodo.21414390

### Competing Interest Statement

The authors have declared no competing interest.

